# Phylogenetic diversity and regionalization in the temperate arid zone

**DOI:** 10.1101/2023.11.01.565216

**Authors:** Ryan A. Folk, Aliasghar A. Maassoumi, Carolina M. Siniscalchi, Heather R. Kates, Douglas E. Soltis, Pamela S. Soltis, Michael B. Belitz, Robert P. Guralnick

## Abstract

*Astragalus* (Fabaceae) is astoundingly diverse in temperate, cold arid regions of Earth, positioning this group as a model clade for investigating the distribution of plant diversity in the face of climatic challenge. Here we identify the spatial distribution of diversity and endemism in *Astragalus*, using species distribution models for 752 species and a phylogenetic tree comprising 847 species. We integrated these to map centers of species richness (SR) and relative phylogenetic diversity (RPD), and used grid cell randomizations to investigate centers of endemism. We also used clustering methods to identify phylogenetic regionalizations. We then assembled predictor variables of current climate conditions to test environmental factors predicting these phylogenetic diversity results, especially temperature and precipitation seasonality.

We find that SR centers are distributed globally at temperate middle latitudes in arid regions, but the Mediterranean Basin is the most important center of RPD. Endemism centers also occur globally, but Iran represents a key endemic area with a concentration of both paleo- and neoendemism. Phylogenetic regionalization recovered an east-west gradient in Eurasia and an amphitropical disjunction across North and South America; American phyloregions are overall most closely related to east and central Asia. SR, RPD, and lineage turnover are driven mostly by precipitation and seasonality, but endemism is driven primarily by diurnal temperature variation. Endemism and regionalization results point to western Asia and especially Iran as a biogeographic gateway between Europe and Asia. RPD and endemism highlight the importance of temperature and drought stress in determining plant diversity and endemism centers.

## INTRODUCTION

The story of angiosperm diversification can be seen from the point of view of margins: how plants left their ancestral tropical origins and gained adaptations to confront the stresses of an increasingly challenging and dynamic climate (Edwards et al., 2010; Arakaki et al., 2011; Folk et al., 2019, 2020; Soltis et al., 2019). Empirical investigations of the diversification process have underlined the difficulty of identifying straightforward causal factors behind such evolutionary radiations (Singhal et al., 2022; Blaimer et al., 2023), but studies of model clades with well-developed biodiversity data offer a clear path for identifying the key underlying determinants of current plant biogeography, diversification, and community composition. As argued by Cavender-Bares (2019), investigations of phylogenetic diversity at low phylogenetic levels are particularly valuable for generating such insights. The two key advantages mentioned are (1) recent radiations have far less risk of being obscured by evolutionary time; and (2) radiations of a tractable size, particularly those with a history of research, can offer rich and diverse data sources that can be further synthesized to derive multifaceted understanding of patterns and processes determining plant diversity (see also Liow et al., 2022).

One of the most obvious and central benefits of clade-based spatial phylogenetics is that it facilitates the study of drivers of diversity in the context of ecological specialization. Investigations of all seed plants or all vascular plants (Thornhill et al., 2016, 2017; Scherson et al., 2017; Allen et al., 2019; Mishler et al., 2020) can elucidate the broadest factors responsible for floristic diversity, but may largely reflect the most diverse and spatially extensive areas of Earth. This approach limits investigation of the margins of plant diversity where evolutionary processes may be active, rapid, and subject to unique environmental drivers (Folk et al., 2020). Clades that specialize in challenging habitats and have recently radiated in those habitats may provide the best case for understanding processes that allow lineages to diversify and thrive at the ecological margins.

*Astragalus* (Fabaceae) is astoundingly diverse in temperate, cold arid regions of Earth (as reviewed in Folk et al., 2020). Morphologically fairly homogeneous and diverging no earlier than the late Miocene, it comprises at least 3,100 species and arose from some of the highest diversification rates documented so far in angiosperms (Folk et al., 2023a). *Astragalus* represents a major radiation in arid extratropical areas of the Northern Hemisphere, often the most diverse and dominant legume in such environments. The colonization of different geographic areas was closely related to major shifts in niche, reflecting different temperature, precipitation, and soil stressors within the arid zone (Folk et al., 2023a). *Astragalus* therefore represents an excellent opportunity to study spatial factors associated with diversification in the face of challenging environments.

*Astragalus* has been the focus of one study on spatial diversity. Maassoumi and Ashouri (2022) used taxonomic and curated distributional records from *Astragalus of World* (https://astragalusofworld.com/) at the level of political units to investigate centers of species richness and endemism, considering the Eastern Hemisphere portion of its range. This distributional area, and more specifically central Asia, likely corresponds to the geographic origin of *Astragalus* and all major clades but one Folk et al. (2023a) and contains much of its species richness. The primary results of Maassoumi and Ashouri (2022) indicated four key hotspots of both species richness and endemism (east to west): Afghanistan, Iran, the Caucasus, and Anatolia. Using Maassoumi and Ashouri (2022) as a starting point, we (1) added the additional context of Western Hemisphere centers of diversity, (2) added a phylogenetic component to the previous endemism analysis, allowing us to distinguish between paleo- and neoendemism, and (3) used gridded occurrence records and niche models, which complement important limitations of records based on political units and allow for finer spatial resolution.

Here, we map measures of species richness, phylogenetic diversity, and endemism globally for *Astragalus.* We first (1) identify hotspots on the basis of species richness and (2) ask if these richest areas are also places of high phylogenetic diversity and neo- or paleoendemism. We then (3) use phylogenetic beta analyses to identify major phyloregions and investigate how abiotic niche shapes lineage turnover between these phyloregions. In a previous study (Folk et al., 2023a), we identified phylogenetically conserved ecological shifts associated with precipitation and temperature variability. Here, in a spatial rather than a phylogenetic context, we identify the environmental factors that best predict diversity and endemism in *Astragalus* and test the importance of the earlier identified biogeographic factors, especially aridity and diurnal temperature variation, for predicting centers of diversity.

## METHODS

### Niche modeling

We use a niche modeling approach implemented in R with manual curation steps to estimate the geographic distribution of individual species of *Astragalus*. Occurrence records were gathered from GBIF (GBIF 2020) and iDigBio as reported previously (Folk et al., 2023a). For each species with at least five records, the accessible area of the niche model was defined by calculating an alpha hull around the cleaned occurrence records of the species (Rabosky et al., 2016). The accessible area was then buffered by the larger value of either 75 km or the 80th percentile of distances between nearest occurrence record pairs. Next, we compiled a predictor set of model covariates including 18 environmental variables, consisting of 13 bioclimatic variables (Bio1, Bio2, Bio4, Bio5, Bio6, Bio8, Bio9, Bio12, Bio13, Bio14, Bio15, Bio16, Bio17), 3 soil variables (nitrogen (cg/kg), sand (g/kg), and soil organic carbon stock (t/ha), and 2 topographic layers (elevation and ruggedness). The bioclimatic variables were chosen among the WorldClim variable set (Fick and Hijmans, 2017) to represent a reduced list of biologically important predictor variables. The soil variables were the average values across the depths of 0–5 cm, 5–15 cm, and 15–30 cm and were provisioned from the International Soil Reference and Information Center (Batjes et al., 2017). The topography layers were provided by EarthEnv (Amatulli et al., 2018). All variables were resampled using bilinear interpolation to a grid cell size of approximately 18 by 18 km following previous methods (Folk et al. 2023).

For each species, we fit an initial Maxent model using cleaned occurrence records as presence points fit to the full predictor set using default Maxent settings. Next, the variance inflation factors (VIFs) of each predictor variable of the initial model were calculated, and predictors were sequentially removed from iteratively fit models until the model did not have a predictor with VIF > 5. This procedure therefore yields unique predictor sets for each species. Once the predictor set was finalized, we optimized the tuning parameters of Maxent using multiple combinations of regularization multipliers and feature classes (see Folk et al., 2023b for specific tuning parameter combinations) using the R package ENMeval (Muscarella et al., 2014). Occurrence and background points were split into training and testing data using block partitioning. In most cases, the model with the lowest AICc value was selected as the top model, but in the few cases where a model had a training and validation area under the curve (AUC) value less than 0.7, we selected the model with the highest validation AUC as the top model. Top models were converted to predicted presence/absence maps using the 10% least presence threshold approach, where the lowest 10% of predicted presence values are deemed absences. We used this threshold because it reasonably balances omission and commission errors and especially avoids over-commission in modeled range estimates.

### Phylogenetic estimate

The sequence data here derive from the NitFix sequencing initiative (Kates et al., 2022), which was based on sequence capture data targeting 100 phylogenetic loci as described in Folk et al. (2021a) and Kates et al. (2023) and comprised 847 species of *Astragalus* (∼27% of currently recognized species). The tree used in spatial phylogenetic statistics has been previously reported (Folk et al., 2023a) and was based on assemblies of NitFix data performed in HybPiper (Johnson et al., 2016) and coalescent analysis in ASTRAL (Zhang et al., 2018), followed by curational steps excluding poorly sequenced taxa and by secondary constraint-based time calibration as reported in Folk et al. (2023a).

### Metrics of diversity

To calculate diversity metrics, we imported the presence/absence matrix derived from distribution models discussed above into Biodiverse v. 3.1 (Laffan et al., 2010). The two focal metrics were species richness (SR) and relative phylogenetic diversity (RPD, i.e., the sum of branch lengths in a grid cell divided by the sum calculated on a tree with no branch length structure; see Mishler et al., 2014). Raw phylogenetic diversity (PD) values have the disadvantage that they are tightly correlated to SR (because PD scales with the number of branches, which scales with the number of species) and therefore are of limited use for identifying diversity independent of SR. RPD, by contrast, contains branch sums in the numerator and denominator and therefore is effectively corrected for species number and does not directly scale with SR. Compared to PD (the direct sum of all phylogenetic branches connecting members of a grid cell), low RPD thus can be interpreted as representing shorter branches than a phylogenetically unstructured community, and high RPD represents longer branches, a property that facilitates generating null distributions via randomization.

Randomizations in Biodiverse were used to investigate whether RPD and phylogenetic endemism are significantly different from the null expectation of phylogenetically unstructured grid cells. We calculated 100 replicates with species randomly assigned to grid cells while maintaining species range size and grid cell SR. From these randomized grid cells, significance values were calculated for RPD; two-tailed *p-*values were calculated directly from the randomizations as reported previously (Folk et al., 2023b). We then calculated relative phylogenetic endemism (RPE; see Mishler et al., 2014) to enable randomizations identifying areas of endemism using the CANAPE method (Mishler et al., 2014). Based on a hierarchical *p-* value procedure using the randomizations (details summarized in Folk et al., 2023b), CANAPE identifies grid cells having a high proportion of neoendemics (small-range taxa with shorter branch lengths than the null expectation), paleoendemics (small-range taxa with longer branch lengths than the null expectation), and mixed endemism (both longer and shorter branch lengths than the null; i.e., a mixture of both neo- and paleoendemics).

### Phylogenetic regionalization

We also used Biodiverse to implement distance measures of phylogenetic turnover and subsequent hierarchical clustering to quantify areas with similar phylogenetic composition, i.e., “phyloregions” (Daru et al., 2017). We then compared these phyloregions to regionalizations that have been recognized previously in broader groups of plants using similar methods. We were particularly interested in the geographic breaks that a regionalization analysis would recover in central Asia, previously recognized as an important area for interpreting the biogeography of *Astragalus* and therefore likely to be a key area for lineage turnover. The metric of turnover was based on the branch lengths of shared and unshared community members as weighted by range size, which therefore downweights the influence of wide-ranging species (Laffan et al., 2016) that may be expected to contain less regionalization information. We manually identified breaks in the hierarchical clusters that delimited well-defined groups of grid cells.

### Models for predicting diversity

We implemented a series of generalized linear models with environmental characteristics as predictors and diversity metrics as the response to understand which environmental factors are best associated with diversity. Models were fit for three responses: SR, RPD, and CANAPE significance categories. The list of predictors included: UNEP aridity index (reported in Folk et al., 2023b), mean annual temperature (Bio1), isothermality (Bio3; a measure of diurnal temperature range, following the arguments of Folk et al., 2023a), temperature seasonality (Bio4), temperature annual range (Bio7; included as another measure of temperature seasonality), annual precipitation (Bio12), temperature of the driest quarter (Bio17; included as a measure of precipitation seasonality), elevation, and three soil predictors to capture proxies for soil fertility (sourced from Hengl et al., 2017): nitrogen content, soil pH, and soil carbon content, and a further two soil predictors that describe physical structure: sand percent and coarse fragment percent. Finally, two variables from Tuanmu and Jetz (2014) were included as measures of forest cover: deciduous forest land cover and coniferous forest land cover. All predictor variables were mean centered and scaled to appropriately compare model coefficient values.

Model selection was performed by AIC and comprised two distinct steps chosen on the basis of the large number of predictors. First, VIFs were calculated on a full model including all 15 environmental variables to control collinearity. The variable displaying the highest VIF was eliminated, and VIFs were recalculated; this procedure was repeated to yield all VIFs < 5. This result is referred to as the “full” model. To further investigate the effect of predictor numbers beyond collinearity alone, standardized coefficients were extracted from the full models, and the top five were used to fit a simpler model, referred to hereafter as the “reduced” model. For each response, three model classes were implemented: (1) a standard linear model (logit model for the categorical CANAPE response) was fit for the full and reduced environmental predictors; this was followed by (2) a linear mixed model for full and reduced predictor sets, with the environmental factors as fixed effects and latitude and longitude as random effects. The purpose of the linear mixed model approach was to identify whether environment-diversity relationships were confounded by spatial autocorrelation by separately partitioning latitudinal and longitudinal variation.

We used a model-based ordination approach for investigating environmental factors that are associated with phyloregions and therefore lineage turnover. The recovery of distinct non-overlapping phyloregions would indicate an association with environment, with coefficients on ordination axes indicating the most important environmental factors distinguishing phyloregions. This issue was addressed using a linear discriminant analysis, a classification approach to identifying factors that best explain group separation, and used the full model predictor set as described above.

## RESULTS

### Species richness

Based on SR discontinuities in the 752 mapped species, seven hotspot regions are recognized here. Proceeding west to east, the largest in extent was western North America (hotspot 1, Fig. 1), an area comprising most of the Great Basin and the southern-central Rockies. Next was Iberia (hotspot 2), specifically the northeast area centered on the Sistema Ibérico, with adjacent richness also in the Alps and western North Africa. Next are regions centered on the Levant (hotspot 3) and Caucasus-Irano-Touran (hotspot 4). Two major hotspot regions in central Asia are a region comprising the Pamirs south to the Hindukush (east Tajikistan to Kashmir; hotspot 5), and the Altai and Sayan mountains west to Lake Baikal (approximately Altai Territory to Irkutsk Oblast, Russia; hotspot 6, which was the largest Eurasian hotspot). Finally, hotspot 7 occurred in central China, approximately from the Hengduans north to Qinghai and Gansu provinces; this is hereafter called the QTP (Qinghai-Tibetan Plateau) region as its boundaries closely correspond to the QTP.

**Fig. 1.**
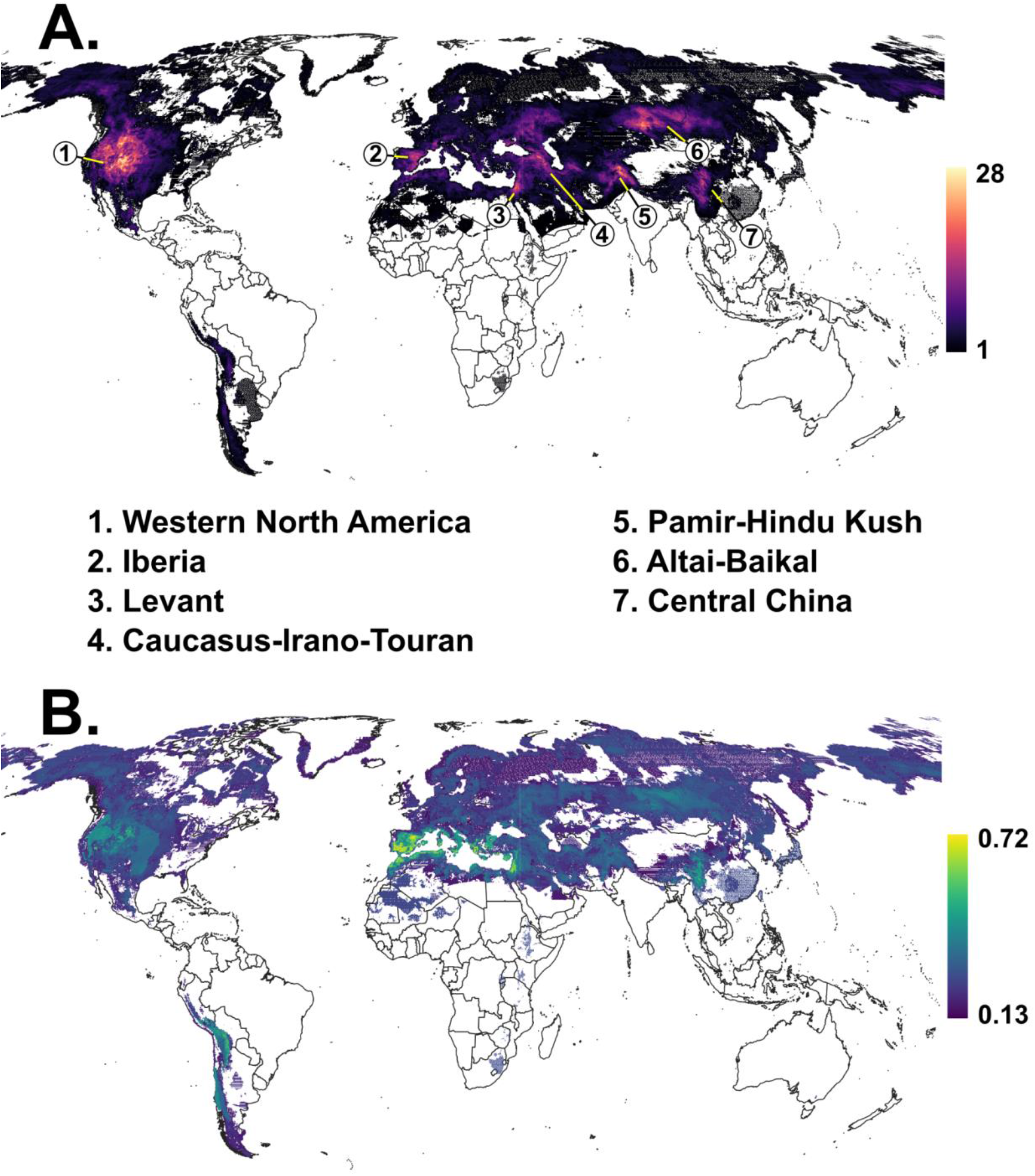
Basic diversity in *Astragalus*. A. Global species richness (SR) of *Astragalus*, with warm colors representing a greater number of species and white areas of land indicating no mapped species. B. Global relative phylogenetic diversity (RPD), with warmer yellow colors indicating higher RPD. Projection: World Robinson (EPSG: 54030).

Of the remainder of the distribution of *Astragalus*, montane and temperate South America and montane East Africa are important in that they comprise the only presence of *Astragalus* in the Southern Hemisphere, but these areas display low species richness. Similarly, the boreal area is extensively colonized by *Astragalus*, but with very low species richness. *Astragalus* has only marginal presence in temperate forest regions, leading to its low presence in eastern North America and East Asia (but not northwestern Europe), and it is completely absent from the global lowland tropics.

### Investigation of sampling bias

Because of the size of *Astragalus* and the approach employed here using incomplete sampling, we investigated potential spatial bias resulting from the use of documented occurrences and species distribution models. We used taxonomically complete country-level checklist data as reported in Maassoumi (2020) and Maassoumi and Ashouri (2022) to investigate whether incomplete sampling influences diversity mapping. There was a clear relationship between species richness estimates based on the occurrence data included in this study and published checklist data (Fig. 2); most countries have between a quarter and half of species sampled, and thus are similar to the overall species sampling. While sampling was variable among countries, the relationship between species richness estimates by checklists and occurrence data was quite strong (linear model, y and x log_10_ transformed: *R*^2^ = 0.7644; *p* < 2.2e-16, linear model *F-*test), with estimated slope 1.0218, therefore close to the 1:1 line which would indicate complete proportionality.

**Fig. 2.**
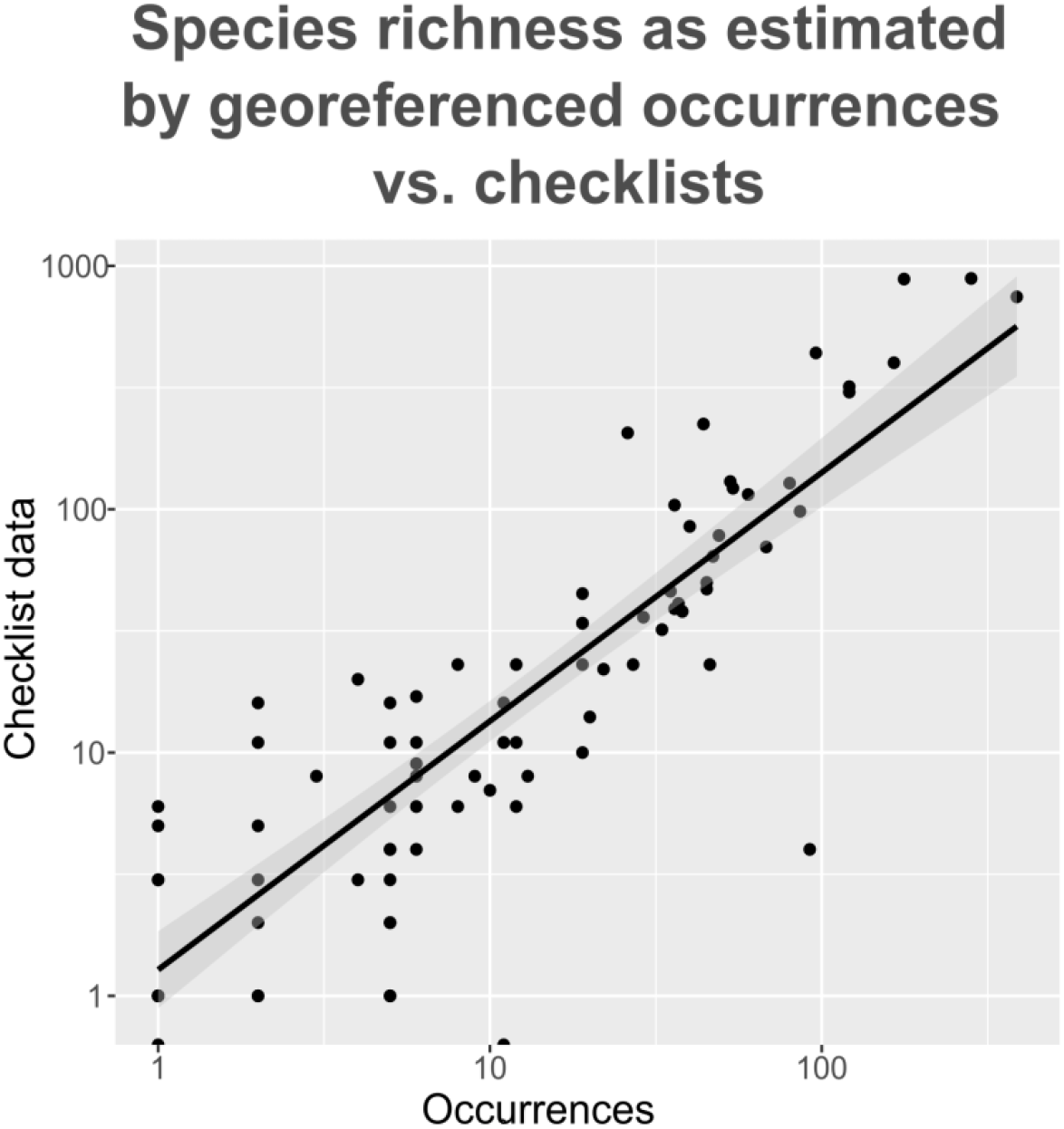
Species richness as estimated by georeferenced occurrence data vs. checklist data. The trend line represents a linear model with a 95% confidence interval (gray curve); countries above the line are undersampled (i.e., more checklist species than expected, particularly on the right tail representing the most diverse countries) compared to the whole study while those below the line are oversampled (fewer checklist species than expected). These deviations were interpreted as minimal. The data were approximately exponentially distributed so both axes are shown as log_10_.

### Relative phylogenetic diversity

Integrating species distribution models with phylogenetic data yielded an overlap of 637 species; this dataset was used for further downstream analysis. Coverage of the phylogeny was incomplete due to excluding species with < 5 records which would be too few for a robust distribution model. RPD showed only moderate alignment with centers of species richness. The highest area on Earth for RPD is Iberia (Fig. 1b), a region larger in extent than the Iberian Hotspot (2); this was part of a larger high-RPD area that comprised the entire Mediterranean Basin including the Levant Hotspot (3). Three further areas showed moderately high RPD: western North America, the QTP region, and the Altai region. The central Andes comprised a final high-RPD region that did not align at all with centers of species richness.

Randomizations of RPD (Fig. 3a) demonstrated greater alignment with species richness hotspots. The Mediterranean Basin is mostly higher in RPD than expected by chance, as are the Andes. However, most of Central Asia is characterized by lower RPD than the null expectation despite having relatively high raw RPD, and western North America is divided into different significance categories, with the Great Basin having significantly lower than expected RPD but the Sierra Nevada and adjacent mountains significantly higher. Two further regions distinguished by greater RPD than expected (but very low species richness) are the highlands of eastern South Africa and temperate East Asia.

**Fig. 3.**
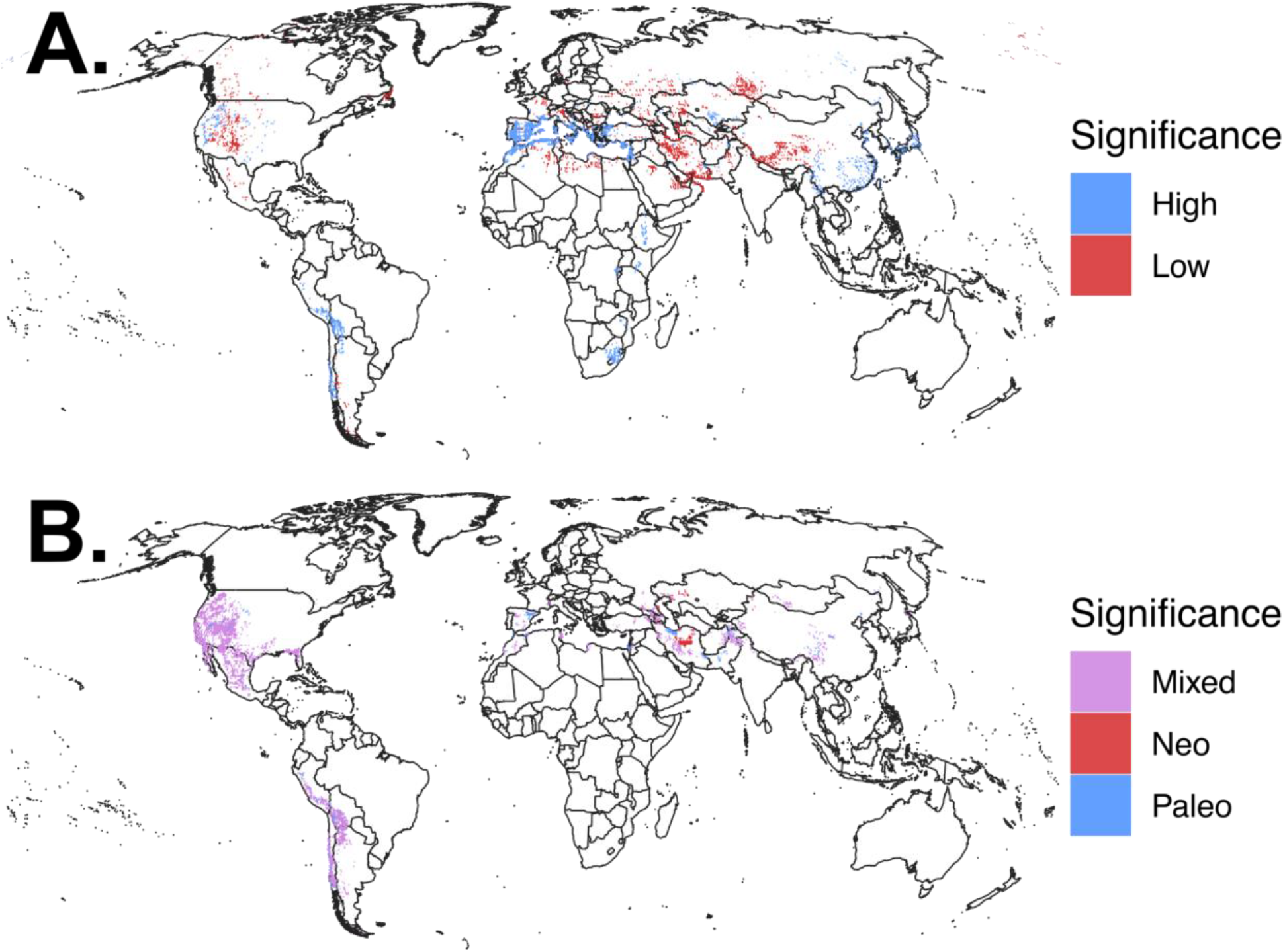
(A) RPD randomization. (B) CANAPE endemism analysis.

### Endemism

CANAPE analysis distinguished large centers of mixed endemism in western North America and the central Andes (hotspot 1; Fig. 3b). A third region of mixed endemism in the Americas occurred along the Southeastern Coastal Plain of North America, an area that is species-poor but with several narrow endemics. Eurasia had more spatially restricted endemism, including the only two paleoendemism centers detected globally: the Pyrenees of Spain (hotspot 2) and the Elburz Mountains of northern Iran. The only globally recognizable area of neoendemism was also in Iran: the Kavir Desert of the north-central region, which was not recognized as distinct in the other analyses. Aside from these areas of paleo- and neoendemism, montane central Iran, the QTP region (hotspot 7), the Caucasus-Irano-Touran (hotspot 4), and the Hindu Kush (hotspot 5) were small areas of mixed endemism.

### Phylogenetic regionalization

The twelve regions recovered by the phyloregionalization analysis (Fig. 4) partly corresponded with recognized SR hotspots (Fig. 1a). In general, the correspondence between regionalization and SR hotspots was less in the Americas than in Eurasia (below) because the three phyloregions in the western United States, while clearly demarcated according to phylogenetic composition distances, are spatially abutted against one another with no SR discontinuity.

**Fig. 4.**
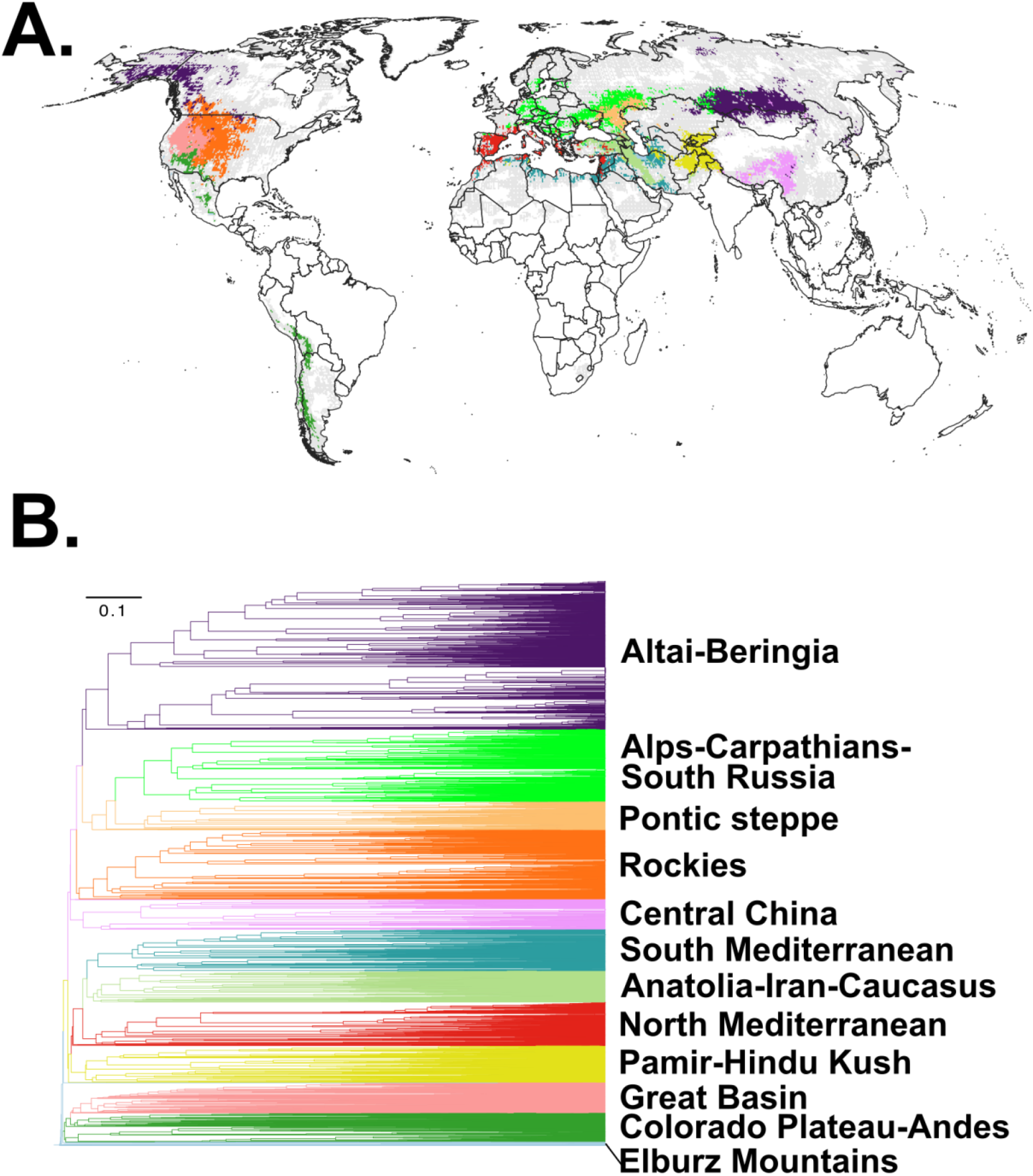
Phyloregionalization. (A) Map of recovered clusters; gray pixels correspond to grid cells with three or fewer species, which were not included in the analysis. (B) Dendrogram of the same grid cells as those plotted in (A), with corresponding colors.

Proceeding from west to east, North America (primarily occupied by the Neo-Astragalus clade) was divided into four regions (Fig. 4a). The northernmost region of Alaska and adjacent Pacific Canada (which was not a SR hotspot; Fig. 4a, dark purple, marked “Altai-Beringia”) formed part of a region with the Altai-upper Mongolia (hotspot 6) and thus was the only connection between the Eastern and Western Hemispheres. The center of western North America was divided into three regions, one corresponding closely to the Great Basin (hotspot 1; Fig. 4a, salmon pink), another corresponding to the Rockies and adjacent plains (also hotspot 1; Fig. 4a, dark orange), and finally a region corresponding to the southern Colorado Plateau and Basin and Range physiographic provinces (Fig. 4a, dark green, marked “Colorado Plateau-Andes”). This last phyloregion also included occurrences in the Sierra Madre of Mexico and Andean South America, therefore marking *Astragalus* in this region as a typical amphitropical disjunction (Raven, 1963; Scherson et al., 2008; Simpson et al., 2017). To understand whether regionalizations were associated with abiotic niche differentiation, we performed a linear discriminant analysis. This analysis (Fig. 5a) identified the Altai-Beringia area as occupying a distinct niche space, whereas the other phyloregions showed considerable overlap.

**Fig. 5.**
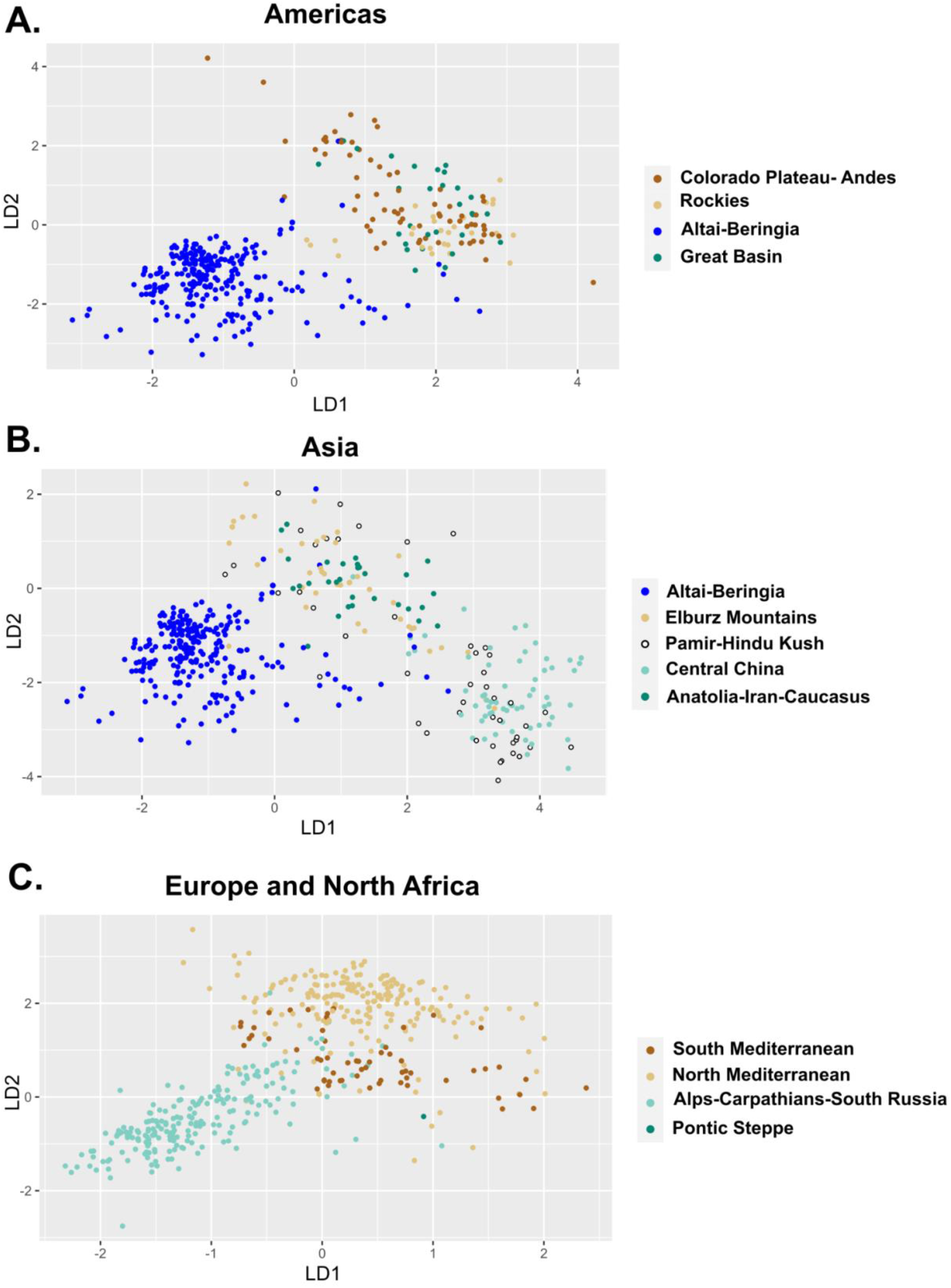
Linear discriminant analysis of the regionalization. This ordination quantifies the relationship between phyloregions and climatic data. A global analysis was conducted but is plotted here by continent for clarity. As noted in the main text, most phyloregions in Asia and the Americas overlap except for the Altai-Beringia region in Asia and North America, the Alps-Carpathians-South Russia region in northern Europe, and to a lesser extent the North and South Mediterranean regions.

Based on the dendrogram (Fig. 4b), the Great Basin and Colorado Plateau-Andes regions were closely related but highly distinct from the other two regions, and not closely related to any Eastern Hemisphere regions. The remaining two regions of North America were more closely related in species composition to regionalizations recognized in Eurasia: the Altai-Beringia region was most similar in phylogenetic composition to the two central-eastern European regions discussed below, and the Rockies region was most similar to the three foregoing regions.

In Europe, three phyloregions were recognized. The North Mediterranean Basin (including but larger than hotspot 2; Fig. 4a, red) was recognized as a phyloregion that also included most of Iberia to the Pyrenees; most of the southern and eastern Mediterranean Basin was a separate South Mediterranean region (discussed below). Much of montane central Europe, from the Alps to the Carpathians and extending west to most of eastern Ukraine and part of southeastern Russia, terminating at the Urals, formed a second phyloregion, the most spatially extensive region in Europe. About half of southwestern Russia was in the third European region, in the general area of the Pontic Steppe and corresponding closely to the drainage basin of the Volga (Fig. 4a, light orange). The linear discriminant analysis of phyloregions (Fig. 5c) showed that all phyloregions that occur in Europe occupy distinct areas of climate space.

In western Asia, the eastern Mediterranean Basin was recovered as a distinct phyloregion (including but larger than hotspot 3; Fig. 4a, blue-green) that also included most of the South Mediterranean Basin in North Africa (sporadically west to the Anti-Atlas; most of the Tell Atlas and Rif west of Tunisia were instead assigned to the North Mediterranean Basin region). This South Mediterranean region also included sporadic areas further east including most of Syria (the Aleppo Plateau was instead assigned to the North Mediterranean) and disjunct northern and southern portions of Iraq and Iran. The most spatially extensive phyloregion in western Asia extended through the Pontic Mountains of the northern half of Anatolia, the Caucasus, and the Zagros Mountains of the western half of Iran (including but larger than hotspot 4; Fig. 4a, olive green). Finally, the most distinctive phylogenetic assemblage of *Astragalus* on Earth (light blue in Fig. 4a; see dendrogram in Fig. 4b) was the Elburz Mountains of northern Iran, which was also one of the two paleoendemism centers. Aside from the Elburz Mountains, all western Asian phyloregions were closely related and shared phylogenetic composition with the North Mediterranean Basin (Fig. 4b).

Considering the rest of Asia, only one phyloregion was recognized in central Asia (hotspot 5 in the Pamirs-Hindu Kush, yellow in Fig. 4a) and two in East Asia, one in the QTP region (hotspot 7; pink in Fig. 4a) and the other in the Altai-upper Mongolia region shared with Beringia as noted above (including but larger than hotspot 6; Fig. 4a, dark purple, labeled “Altai-Beringia”). The Pamir-Hindu Kush phyloregion was similar in phylogenetic composition to western Asia, whereas the QTP region, much like the Altai-Beringia, had greater affinities to western North America. Similar to the Americas, a linear discriminant analysis (Fig. 5b) of the Asian phyloregions showed overlap except for the highly distinct Altai-Beringia region.

### Modeling

For all three responses (SR, RPD, and CANAPE significance), inclusion of latitude and longitude as random effects was strongly favored (ΔAIC >> 100) as was the inclusion of the fixed effects (that is, the environmental variables; ΔAIC >> 100; see Supplementary Table S1). Variable choice differed across the models, but aridity and Bio7 (temperature annual range) were eliminated from all models due to high collinearity in this dataset. Full model results including normalized fixed effects and model choice statistics are available in Supplementary Tables S1 and S2.

For SR, a full model including 11 predictors was favored; the most important predictor as measured by standardized coefficients was Bio12 (mean annual precipitation; negative relationship), closely followed by elevation (positive relationship), befitting the close association of species richness in *Astragalus* with high elevation and low rainfall. Model fit for the RPD response favored a full model with ten predictors, and demonstrated that Bio17 (precipitation of driest quarter; negative relationship) and Bio1 (mean annual temperature; positive relationship) were most important with near-identical coefficients; these predictors were likewise expected and reflect the importance of the Mediterranean Basin in particular for *Astragalus* RPD. Mean annual temperature showed particularly high separation among the RPD significance categories (Fig. 6e), with low RPD (short branches) associated with lower temperature. RPD significance was also plotted against isothermality (although not in the best model, this was done to compare with the CANAPE analysis below), demonstrating an association of long branches with higher diurnal temperature change. Finally, the favored model for the CANAPE significance response was a reduced model with five predictors, with Bio3 (isothermality) by far the most important predictor. All significant endemism categories were associated with increased isothermality (greater diurnal temperature contrast), especially mixed and paleoendemism (Fig. 6g). Precipitation of the dry season (Fig. 6h) also showed a strong separation among endemism categories, with significance associated with lower dry season precipitation, an association that was most pronounced for neoendemism.

**Fig. 6.**
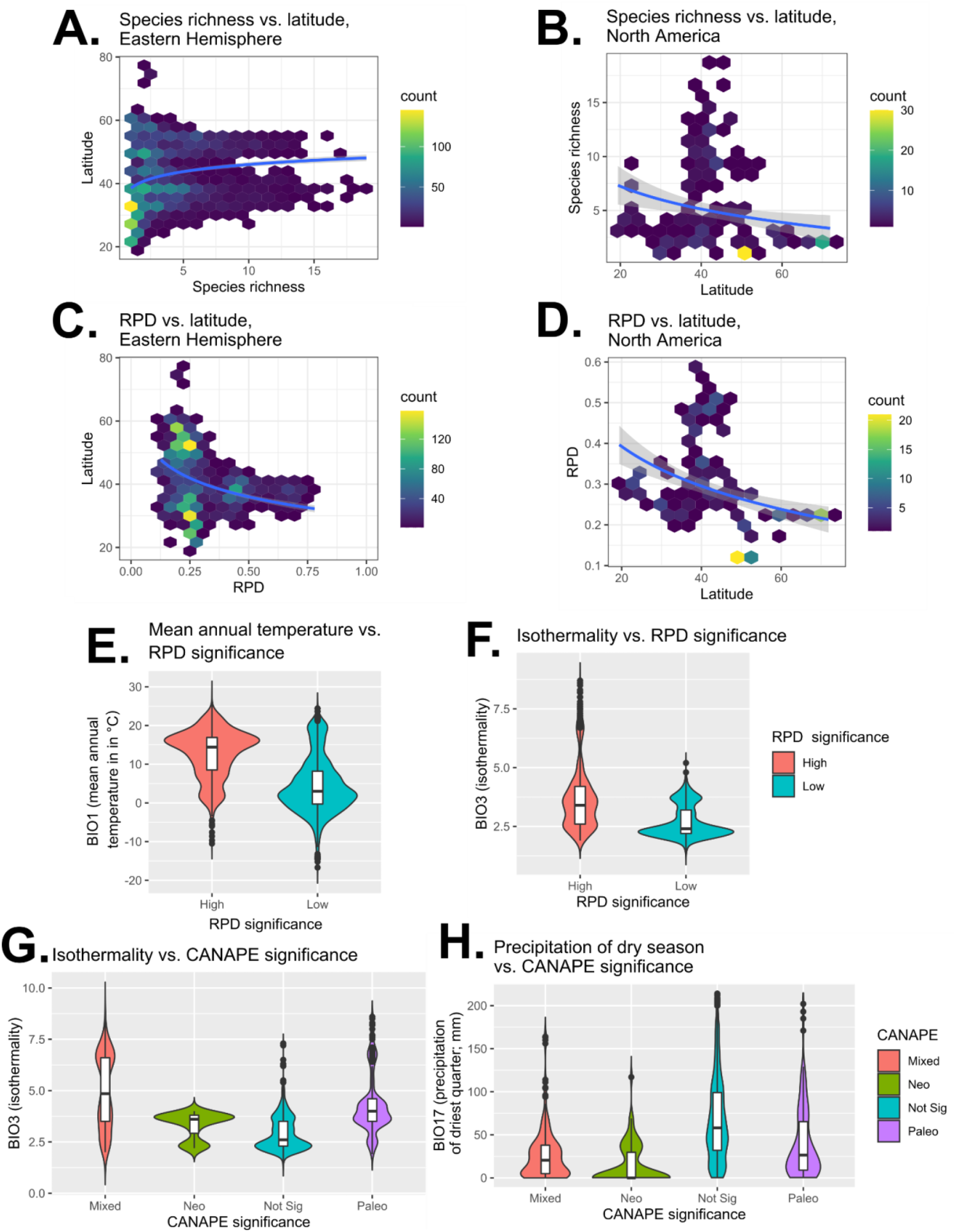
Diversity-environment relationships. A–B: S vs. latitude. C–D: RPD vs. latitude. A and C show Eurasia; B and D show North America. Because diversity statistics showed an approximately exponential relationship with latitude, trend lines (blue) and 95% confidence intervals (gray) represent logarithmic models. E–F: Mean annual temperature (E) and isothermality (F) vs. RPD significance categories. G–H: Isothermality (G) and precipitation of the dry season (H) vs. CANAPE significance categories.

Plots investigating latitudinal patterns revealed contrasting relationships between North American and Eurasian species. For North America, both SR (Fig. 6b) and RPD (Fig. 6d) show a typical latitudinal response, increasing towards the equator. Eurasian species demonstrate partially consistent and partially contrasting relationships, with RPD showing a typical increase towards the equator but SR slightly increasing towards the poles.

## DISCUSSION

### *Hotspots of* Astragalus *diversity*

We identified key hotspots of SR for temperate arid specialists in *Astragalus*, recognizing seven hotspots in total. These occur exclusively in the Northern Hemisphere, closely follow the limits of major montane physiographic regions, and span the continental distribution of the genus. North America, not previously examined in spatial detail for *Astragalus*, comprises a single large hotspot covering much of montane western North America. The Eurasian hotspots as recognized here show a close correspondence with hotspots reported in Maassoumi and Ashouri (2022), but with important interpretational differences. Maassoumi and Ashouri primarily relied on country-level occurrences, and most analyses did not correct for the differing areas of political units, creating different emphases than would be yielded by grid-cell statistics. Our use of a grid-cell approach for identifying hotspots allows for finer spatial distinctions. Notably, based on grid cells and the sampling achieved here, Anatolia and the portions of Iran independent of the Caucasus are not recognized as independent hotspots (although these regions were distinct in randomization and regionalization analyses). This analysis also identifies the Chinese hotspot as primarily north of (not in) the Himalayas, does not recognize the Mediterranean Basin in its entirety (only the Levant, but see RPD below), and more finely delimits several central Asian hotspots.

Beyond the hotspots recognized here, the distribution of *Astragalus* extends into the Southern Hemisphere in montane South America and East Africa and into boreal regions. These additional areas of distribution are very low in SR; in boreal areas the ecological role of *Astragalus* is primarily replaced by the close relative *Oxytropis* (Meyers et al., 2013; Kholina et al., 2016), and no close relatives occur in the Southern Hemisphere, where similar habitats would be occupied by *Lupinus* and other unrelated legumes. As the correlation of the results reported here against checklist data in Maassoumi (2020) demonstrates, our hotspot results are robust to incomplete sampling.

### Alignment of SR and RPD

Our second major goal was to assess the alignment of SR hotspots with RPD. The rationale behind assessing SR-RPD mismatch was to determine areas of Earth that have particularly old or young lineages. RPD values that scale approximately with SR indicate that phylogenetic assemblages of species are within expectations; when RPD is high in comparison to SR this would indicate a surfeit of ancient diversity, but a low RPD compared to SR would indicate a profusion of recent lineages. As well as comparing raw values, we further tested our expectations through null randomizations.

We identified primary areas of the distribution of *Astragalus* where RPD contrasts with SR: the Mediterranean Basin, western North America, and central-eastern Eurasia (the QTP region to the Altai-Baikal region). Among these, the Mediterranean Basin stands out as containing the highest *Astragalus* RPD on Earth. Randomizations revealed that, despite high RPD overall, this area contains a north-south division in RPD, with the northern half of the Mediterranean and the adjacent eastern and western extremes of the South Mediterranean (specifically, the Levant and the Atlas mountains) having higher RPD than expected. Thus, this area is richest in ancient phylogenetic lineages. The remaining South Mediterranean Basin, comprising the Tell Atlas and Anti-Atlas ranges, is lower than null RPD expectations, suggesting this area is instead rich in younger lineages and possibly the site of recent diversification. The north-south RPD pattern recovered in the Mediterranean Basin very closely reflects phylogenetic regionalization (see below), suggesting the pattern is driven by distinct lineages along the north-south gradient. This is the first detailed spatial phylogenetic analysis we are aware of covering the whole Mediterranean Basin (see Cheikh Albassatneh et al., 2021), but these results parallel an angiosperm-wide study focusing on the Gibraltar Strait (Molina-Venegas et al., 2015) that found a north-to-south gradient in phylogenetic clustering and turnover. The primarily north-south divide of the Mediterranean Basin has multiple potential explanations, including distinct orogenies, a north-south gradient in aridity and temperature, and separate climatic refugial histories during Pleistocene glaciation. A north-south divide may represent a general pattern of the area ripe for investigation of these mechanisms in other clades.

The additional primary centers of high RPD included western North America, the QTP region, and the Altai-Baikal region. Western North America and China share a strong east-west pattern, with older lineages than expected in the east and younger lineages than expected in the west; both east-west gradients are concordant with those found in recent analyses of Chinese angiosperms (Lu et al., 2018; Hu et al., 2022) and liverworts (Qian et al., 2023), Californian vascular plants (Thornhill et al., 2017), and North American seed plants (Mishler et al., 2020). Both gradients are thought to reflect climatic and geological processes. The east-west gradient in China is thought to reflect the differing ages of flora assembly across China, and particularly the contrast between ancient temperate forests and the relatively recent orogeny of the Qinghai-Tibetan Plateau region and the recent formation of the Asian monsoon (Lu et al., 2018). Similarly, in North America, younger branches than expected are centered in western North America, which is thought to reflect recent radiation in arid western regions. Eastern North America is instead characterized by longer branches than expected, reflecting older forest communities that survived in more climatically stable eastern regions (Mishler et al., 2020). Similarly, two recent spatial phylogenetic analyses of the California flora (Kraft et al., 2010; Thornhill et al., 2017) identified significantly shorter branches primarily in the south and eastern dry areas of California, especially the Great Basin, a result closely associated with relatively recent aridification of the North American interior. The finding of Altai as a center of young lineages has also been recovered in a recent analysis of woody seed plants (Wang et al., 2022), but a wider analysis has shown less consistency (Lu et al., 2018). Overall, the RPD gradients we recovered closely parallel patterns seen in other plant groups and closely follow climatic gradients, especially precipitation (Thornhill et al., 2017).

### Implications of phylogenetic regionalization for the biogeography of the Eurasian interior

Our third goal was to use a phylogenetic regionalization approach to identify patterns in global lineage turnover, focusing particularly on connections between middle Eurasia, the ancestral area of *Astragalus* (Folk et al., 2023a), and other regions of the distribution. Maassoumi and Ashouri (2022) argued that Iran is a biogeographic gateway that links the western and eastern parts of Eurasia and thus serves as a crossroads for diversity that is related to both eastern and western taxa. Our results are consistent with this model, with an additional insight added by the phylogenetic regionalization: this high diversity of Iran reflects disparate evolutionary assemblages, including early-diverging lineages. The presence of multiple phyloregions within a relatively small area reflects Iran as a crossroads, with representatives of early lineages that dispersed from west Asia and colonized Eurasia, Africa, and the Americas (Folk et al., 2023a). The north of Iran is characterized by the most globally distinct lineages in *Astragalus*, with a unique composition of recognized taxonomic sections including *Astragalus* sects. *Cremoceras, Cystium* (endemic), *Hololeuce,* and *Incani*, and an additional three important influences from central Asia (e.g., shared species in *Astragalus* sect. *Erioceras*), the South Mediterranean, and northern Anatolia (e.g., reflected by widely distributed species of *Astragalus* sect. *Cremoceras, Hololeuce,* and *Incani*).

Central to East Asia also proved complex and can be identified as a similar but secondary biogeographic crossroads: the central Asian diversity hotspots (Pamir-Hindu Kush and Altai-Baikal) primarily had Mediterranean and other European lineage affinities, but both north Asia and the QTP region clearly had strong affinity with North America as well, reflecting the Asian affinities of all North American *Astragalus* that are not members of the Neo-Astragalus clade (Folk et al., 2023a). The latter point is reflected in high-latitude tundra species (e.g., *A. eucosmus* and *A. umbellatus*) spanning the two regions and lower-latitude North American species (e.g., *A. adsurgens*, *A. canadensis*, *A. oreganus*) that have close relationships with Asian species. Thus, an east-to-west gradient was recovered across Eurasia, with increasingly shared diversity with the Americas eastward. Within western Eurasia, this study recovers the major phylogenetic turnover occurring across the Mediterranean Basin approximately comprising the northern and southern halves. While the Mediterranean Basin is often thought of as a unit inclusive of North Africa, a recent investigation of phylogenetic turnover in angiosperm trees recovered a similar north-south divide within the European side of the Mediterranean Basin (Cheikh Albassatneh et al., 2021) and across the Strait of Gibraltar in an analysis focusing on the western Mediterranean (Molina-Venegas et al., 2015).

Within North America, phyloregionalizations closely followed major physiographic provinces, namely the Great Basin, Rockies, and Colorado Plateau. The primary novel result was recovering the South American species, long known to have North American phylogenetic affinity, with the Colorado Plateau. *Astragalus* has attracted repeated interest as an archetypal amphitropical disjunction. Previous studies (Scherson et al., 2008; Folk et al., 2023a) have investigated multiple origins of different lineages of South American *Astragalus* species from within the Neo-Astragalus clade, species that dispersed from North America in the early Pleistocene (Folk et al., 2023a). Previous studies did not attempt to discern where in North America these lineages originated, but this study identifies the Sierra Madre and Basin and Range of Mexico and the southwestern United States as the closest plant communities in terms of shared lineages. These results accord with some but not all examples of amphitropical disjunctions. The most recent exhaustive review (Simpson et al., 2017) demonstrates differing distributions rooted in climate between those that are coastal Pacific (e.g., *Fragaria chiloensis*) vs. the interior dry areas of the two continents (e.g., *Larrea tridentata*), where *Astragalus* fits in the interior pattern as reflective of its arid adaptations.

### *Endemism centers of* Astragalus

The most complex endemism area recovered in this analysis corresponds to modern-day Iran. Maassoumi and Ashouri (2022), who originally identified Iran as the key biogeographic province for *Astragalus*, argued for this status primarily on the basis of outstanding endemism in terms of species counts. Results based on phylogenetic endemism in this study were also consistent with Iran’s status as a biogeographic gateway and add to this hypothesis by distinguishing centers of neo- and paleoendemism using a quantitative phylogenetic approach. Iran is distinguished by having extensive mixed endemism as well as the only global neoendemism area and one of the two paleoendemism areas; these regions were closely associated with the major mountain ranges of Iran. Much like the phyloregion analysis (above), Iran’s status as a biogeographic crossroads is underlined by an exceedingly rich *Astragalus* flora that contains early-diverging lineages as well as the products of recent radiation.

Outside of Iran, results for endemism analysis in Eurasia yielded a narrower emphasis compared to the country-level results of Maassoumi and Ashouri (2022). Three further major Eurasian endemism centers were recovered: the mountains of northern Iberia (not recognized in Maassoumi & Ashouri), the Pamir-Hindu Kush area (recognized as “Middle Asia” in Maassoumi & Ashouri), and the QTP region (recognized as part of “China” in Maassoumi & Ashouri). The QTP region as delimited here corresponds very closely to the QTP delimitation of Ye et al. (2019). All of these regions correspond closely to recognized mountainous physiographic provinces, but plant phylogenetic turnover in central Asia is poorly studied overall.

The endemism results for the Americas overall can be compared to a series of papers covering North American seed plants (Mishler et al., 2020) and flowering plants (Earl et al., 2021), and vascular plants of Chile (Scherson et al., 2017) and California (Thornhill et al., 2017). Our results were most similar to those concerning North America (Thornhill et al., 2017; Mishler et al., 2020; Earl et al., 2021): we identified the southwestern US and montane Mexico as primarily associated with mixed endemism, with the interior Great Basin instead showing more neoendemism. *Astragalus* shares its tolerance to dramatic temperature swings with numerous other local radiations in plants (Comstock and Ehleringer, 1992) that reflect rapid diversification in this area. However, while in this study the Andes are primarily centers of mixed endemism, (Scherson et al., 2017) found that the northern Andes contained significant neoendemism. The present analysis focuses on several distantly related and relatively species-poor radiations of the Andes (∼90 species in total; Johnston, 1947; Folk et al., 2023a) and does not include the major evolutionary radiations such as lupines (Hughes and Eastwood, 2006) that reflect unique aspects of rapid Andean diversification (Lagomarsino et al., 2016), thus likely not reflecting recent radiations in the region. That said, the species richness imbalance in *Astragalus* across the Northern and Southern Hemispheres mirrors diversity patterns in other plant and animal groups such as saxifrages and ants (Folk et al., 2021b; Economo et al., 2018) and may reflect differences in land mass or in historical climatic stability, which merit further investigation.

### *Environmental predictors of* Astragalus *endemism and diversity*

Our final goal was to identify factors that shape diversity and regionalization in *Astragalus.* We find that the distribution of diversity in *Astragalus* across multiple measures is shaped by the combination of elevation, temperature variation, and especially precipitation, yet the relative importance of these factors differs when considering different facets of diversity. The finding of isothermality as the most important endemism predictor stands out because this factor is not often studied in plant distribution modeling but it accords well with a recent study demonstrating strong phylogenetic conservatism of diurnal temperature niche in *Astragalus* (Folk et al., 2023a). A key finding of this earlier study was that there is a major biogeographic split between Eurasian *Astragalus* and species of the Americas that correlates strongly with diurnal temperature variation. The present paper adds to these findings by showing that this result relates specifically to endemism: isothermality was a weak predictor of SR and was not included in the RPD model at all (Table S2). Isothermality represents the strength of change in temperature over the course of the day. As discussed by Folk et al. (2023a), this variable represents a major physiological challenge for plants in typically arid regions, requiring morphological and physiological adaptations (Myster and Moe, 1995; Ramsay, 2001; Liu et al., 2013) that are still poorly understood (Folk et al., 2023a). Whether isothermality has a direct effect on diversity, or indirectly affects diversity through aridity or other factors, it is clear that *Astragalus* lineages have become adapted to specialize and thrive in regions with strong diurnal climatic variability.

While isothermality was most important for endemism, this study finds that precipitation factors are most important for diversity overall, with regions of lower precipitation and high elevation most important in the SR model and areas with low precipitation and temperature seasonality in the RPD model. Precipitation, topography, and seasonality, while less associated with delimiting major clades than isothermality (Folk et al., 2023a), are the most significant factors for shaping species diversity hotspots in the dry montane interior of North America and Eurasia.

### Drivers of phylogenetic regionalization

The partitioning of phylogenetic diversity across space is thought to partly reflect environmental factors operating as drivers through phylogenetic niche conservatism (Daru et al., 2017). Thus, major breaks in lineage turnover may correspond to physiological or other environmental constraints that distinguish phylogenetic assemblages. While our understanding of diversity transitions across west and central Asia remains poor, the drivers of lineage turnover have been studied in other plant clades, particularly at global scales. In previous studies (Sun et al., 2014; Scherson et al., 2017; Spalink et al., 2018; Cheikh Albassatneh et al., 2021; Carta et al., 2022), including one previous investigation in legumes (Ringelberg et al., 2023), temperature and precipitation contrasts between regions have been interpreted as closely associated with lineage turnover. We investigated a broad set of factors, finding that rather than overall temperature or precipitation, the primary factors associated with lineage turnover were related to diurnal variation and annual seasonality, especially temperature seasonality. The most important predictors of the linear discriminant first axis by far were, in descending order, temperature seasonality (Bio4), mean annual precipitation (Bio12), and isothermality (Bio3). The second discriminant axis was similarly dominated by three primary predictors, which were temperature seasonality (Bio4; also on axis 1), elevation, and soil coarse fragment percent (see Table S3 for a complete listing of standardized coefficients).

In terms of the spatial distribution of regionalization, the major pattern was a north-to-south divide that spans the Northern Hemisphere. The most distinct transcontinental region, the Altai-Beringia phyloregion, occurred in a highly distinct abiotic niche space in the north, centered on 50°-60° N and distinguished by both discriminant axes (Fig. 5a-b). An approximately equivalent niche space at similarly high latitudes in Europe and western Russia is instead occupied by the Alps-Carpathian-South Russia region (Fig. 5c; note Fig. 5 represents a global analysis with panels A-C showing the same climate space). A secondary north-south divide was recovered between the North and South Mediterranean regions. While the separation was not as dramatic as with the highest-latitude phyloregions (Fig. 5c), the two Mediterranean regions show separation along the second discriminant axis (thus primarily reflecting gradients in temperature seasonality; see also Cheikh Albassatneh et al., 2021).

The remaining mid-latitude phyloregions show broad abiotic overlap both within a continent and especially between continents (compare Fig. 5A-C, all plotted on shared axes). Overall, we interpret this to suggest a north-south divide across all three primary continents containing *Astragalus* species that is driven by abiotic niche differentiation, primarily by gradients in climatic variation centered on harsh, marginal habitats in the Eurasian and North American interior highlands, while the more numerous southern regions and their primarily east-to-west gradient in lineage turnover (discussed above) are not clearly distinguished by climatic, topographic, soil, and vegetation cover factors as measured here.

### Taxonomic interpretation of diversity hotspots

The SR hotspots recognized in this paper partly reflect apparent local radiations of what are currently recognized as taxonomic sections. The Americas (hotspot 1) are dominated by Neo-Astragalus, and a previous study has found extensive non-monophyly of Barneby’s sectional system (Barneby, 1964), so this discussion will focus on recognized lineages in Eurasia which appear to be closer in delimitation to phylogenetic results while still in need of revision (Kazempour Osaloo et al., 2005; Su et al., 2021; Folk et al., 2023a); the taxonomy follows the most recent database of *Astragalus* taxonomy (https://astragalusofworld.com/). Hotspot 2 (Iberia) reflects a large proportion of species from *Astragalus* sects. *Poterion, Chropus, Tragacantha, Alopecuroidei*, and *Astragalus*. These sections correspond to monophyletic groups A and C as recovered by Folk et al. (2023a). Hotspot 3 (the Levant) reflects *Astragalus* sects. *Sesamei, Alopecuroidei, Astragalus, Tragacantha, Poterion, Macrophyllium*, and *Pterophorus*, as well as several of the annual sections (Azani et al., 2017), thus primarily groups A and B of Folk et al. (2023a). Hotspot 4 (Caucasus-Irano-Touran) reflects *Astragalus* sects. *Caprini, Incani, Pterophorus, Malacothrix, Ornithopodium, Hymenostegis, Alopecuroidei, Astragalus, Ornithopodium, Onobrychoidei, Hypoglottidei, Stereothrix, Adiaspastus, Tragacantha, Macrophyllium, Hololeuce, Synochreati, Tricholobus, Glycyphyllos, Rhacophorus, Platonychium*, *Anthylloidei*, *Microphysa*, *Campylanthus, Acanthophace*, and *Oroboidei*; in accordance with its very strong diversity and phylogenetic turnover, many recognized taxa occur in this area (groups A, B, C, and especially F and G of Folk et al. [2023a], which account for most of these sections). Hotspot 5 (Pamir-Hindu Kush) is rich in species from *Astragalus* sects. *Caprini, Aegacantha, Incani, Dissitiflori, Alopecuroidei, Astragalus, Craccina, Galegiformes, Hypoglottidei, Brachycarpus, Komaroviella, Oroboidei, Gontscharoviella, Coluteocarpus, Chlorostachys, Hypsophilus, Lithoon, Pseudoammotrophus, Erionotus, Corethrum, Rechingeriana, Chaetodon, Pendulina*, and *Ammotrophus* (thus groups A, B, H, and especially group F of Folk et al. [2023a], which accounts for more than half of these sections). Hotspot 6 (Altai-Baikal) primarily contains species from *Astragalus* sects. *Dissitiflori, Onobrychoidei, Ornithopodium, Craccina, Uliginosi, Helmia, Hemiphragmium, Trachycercis, Brachycarpus, Oroboidei*, and *Chrysopterus* (thus groups B and F of Folk et al. [2023a], except sect. *Uliginosi* in group C). Finally, hotspot 7 (the QTP region) is rich in species from *Astragalus* sects. *Skythropos, Trachycercis, Brachycarpus, Cenantrum, Komaroviella, Poliothrix, Chryspoterus, Melilotopsis, Tropidolobus, Ebracteolati*, and *Hypsophilus* (distributed approximately equally in groups A, B, D, B, and F of Folk et al. [2023a]). Thus, while smaller delimited taxa were more uniquely distributed in SR hotspots, the higher-level clades show strong overlap among adjacent hotspots and an overall east-to-west pattern of lineage replacement that reflects the phyloregionalization.

### *Missing coverage in* Astragalus *and challenges of sampling large clades*

In this study, we integrated hundreds of species distribution models with the largest *Astragalus* phylogeny constructed to date, both in terms of taxa and genes (Folk et al., 2023a). However, this data assembly must be considered in the backdrop of the logistical challenges of sampling “mega-genera” (Frodin, 2004), artificial taxonomic units that by their very size are resistant to systematic investigation (Folk et al., 2018). Our effort has covered about a quarter of this diversity, so the potential for bias is important to consider. However, while complete sampling would always be preferable, we have several reasons to believe that we present a reasonable estimate of the spatial patterns of diversity. The phylogenetic result has been investigated previously using a comprehensive taxonomic database (Folk et al., 2023a), demonstrating that major clades in *Astragalus* are sampled approximately equally, although the Americas are moderately oversampled. Likewise, as demonstrated here through a correlation analysis against checklist data, while both over- and undersampling are observed, the overall pattern demonstrates that our results are very close to proportionality (Fig. 2). That is, the overall comparison of niche models to checklists reveals a nearly 1:1 relationship between the two, making it unlikely at the global scale that our results are significantly undermined by sampling. However, detailed inferences, particularly in under-digitized areas of central Asia (Meyer et al., 2015; Troudet et al., 2017), could be sensitive to sampling. Overall, the analyses here represent a new framework for investigating biogeography both in *Astragalus* and in the broader central Asia region, generating testable hypotheses that will be amenable to further investigation with increased sampling.

### Conclusions

We identified centers of SR across the Americas and Eurasia in *Astragalus*. RPD centers reflect a subset of these SR regions, with the Mediterranean Basin forming the most important center for lineage diversity. Similar to previous work, Iran and adjacent regions of west Asia form a biogeographic gateway across Eurasia, distinguished by its rich *Astragalus* flora with both eastern and western affinities. The additional diversity centers and phyloregions found here accord with previous work particularly in Asia and North America. Phylogenetic regionalization, the first to be performed in *Astragalus* and apparently the first large-scale phyloregionalization effort in several areas of its distribution, identified an east-to-west gradient across Eurasia, with East Asia sharing significant lineage diversity with the Americas. Within each of these continental regions, a secondary north-south lineage turnover was recovered, driven primarily by seasonality gradients and reflecting phylogenetic assemblages that specialize in harsh habitats subject to sharp temperature and precipitation variation. More broadly, daily and seasonal temperature variability, which are key physiological challenges for plants and closely associated with arid areas, are the most important predictors of diversity and endemism, reflecting the ecological specialization of *Astragalus* in marginal habitats.

## Acknowledgments

This work was supported by the National Science Foundation (USA), grant number DEB-1916632.

---

**Table S1.** Model choice.

| Model | SR AIC | RPD AIC | CANAPE AIC |
| --- | --- | --- | --- |
| GLM, full model | 13446.69 | 6533.545 | 1752.505 |
| GLM, simple model | 13511.2 | 6717.223 | 1759.497 |
| LMM, full model | **12700.66** | **5672.439** | 1436.847 |
| LMM, simple model | 13106.43 | 5846.19 | **1427.321** |
| ΔAIC | 405.77 | 173.751 | 9.526 |

**Table S2.** Model parameters. Boldface indicates the predictor with the highest normalized coefficient; n/a means the predictor was not included in the favored model.

| **Model predictor** | **Best SR model**<br>**normalized**<br>**coefficient** | **Best RPD model**<br>**normalized**<br>**coefficient** | **Best CANAPE**<br>**model**<br>**normalized**<br>**coefficient** |
| --- | --- | --- | --- |
| UNEP aridity index | n/a | n/a | n/a |
| Bioclim 1 | n/a | 0.22168 | n/a |
| Bioclim 3 | 0.28034 | n/a | **-2.0753** |
| Bioclim 4 | n/a | 0.18451 | n/a |
| Bioclim 7 | n/a | n/a | n/a |
| Bioclim 12 | **-0.61164** | n/a | n/a |
| Bioclim 17 | -0.17612 | **-0.22562** | 0.4780 |
| Elevation | 0.58750 | n/a | n/a |
| Soil nitrogen<br>content | 0.10811 | 0.04191 | -0.4006 |
| Soil pH | -0.42719 | -0.15910 | -1.1461 |
| Soil organic carbon<br>content | -0.18769 | -0.01053 | n/a |
| Soil sand content | -0.40522 | -0.18343 | n/a |
| Soil coarse<br>fragment content | 0.19705 | 0.14188 | -0.5289 |
| Landcover –<br>coniferous | 0.18848 | 0.01425 | n/a |
| Landcover –<br>temperate broadleaf | 0.05984 | -0.04017 | n/a |

**Table S3.** Standardized coefficients for the linear discriminant analysis of phyloregions. Boldface: most important predictor (greatest absolute value).

| Predictor | LD1 | LD2 |
| --- | --- | --- |
| UNEP aridity index | -0.5478958 | 0.09480095 |
| Bioclim 1 | -0.2008205 | -0.2812965 |
| Bioclim 3 | -2.3202117 | 0.1525999 |
| Bioclim 4 | **-6.4748148** | **-1.6993533** |
| Bioclim 7 | 0.53783307 | -0.4567682 |
| Bioclim 12 | 4.00296857 | 0.55255882 |
| Bioclim 17 | -0.1193015 | 0.12990309 |
| Elevation | 0.78126727 | -1.5887548 |
| Soil nitrogen content | 0.30779185 | -0.2457603 |
| Soil pH | 0.33867222 | 0.22581823 |
| Soil organic carbon content | 0.19949427 | 0.15285302 |
| Soil sand content | -0.030481 | -0.1300963 |
| Soil coarse fragment content | -0.1055599 | 0.55632184 |
| Landcover – coniferous | 0.14640101 | 0.01618512 |
| Landcover – temperate broadleaf | -0.0836142 | -0.2052969 |

